# Global connectivity of the frontoparietal cognitive control network is related to depression symptoms in the general population

**DOI:** 10.1101/185306

**Authors:** Douglas H. Schultz, Takuya Ito, Levi I. Solomyak, Richard H. Chen, Ravi D. Mill, Alan Anticevic, Michael W. Cole

## Abstract

We all vary in our mental health, even among people not meeting diagnostic criteria for mental illness. Understanding this individual variability may reveal factors driving the risk for mental illness, as well as factors driving sub-clinical problems that still adversely affect quality of life. To better understand the large-scale brain network mechanisms underlying this variability we examined the relationship between mental health symptoms and resting-state functional connectivity patterns in cognitive control systems. One such system is the frontoparietal cognitive control network (FPN). Changes in FPN connectivity may impact mental health by disrupting the ability to regulate symptoms in a goal-directed manner. Here we test the hypothesis that FPN dysconnectivity relates to mental health symptoms even among individuals who do not meet formal diagnostic criteria but may exhibit meaningful symptom variation. We found that depression symptoms severity negatively correlated with between-network global connectivity (BGC) of the FPN. This suggests that decreased connectivity between the FPN and the rest of the brain is related to increased depression symptoms in the general population. These findings complement previous clinical studies to support the hypothesis that global FPN connectivity contributes to the regulation of mental health symptoms across both health and disease.

**AUTHOR SUMMARY:** Understanding how large-scale network interactions in the brain contribute to (or serve a protective role against) mental health symptoms is an important step toward developing more effective mental health treatments. Here we test the hypothesis that cognitive control networks play an important role in mental health by being highly connected to other brain networks and able to serve as a feedback mechanism capable of regulating symptoms in a goal-directed manner. We found that the more well-connected the frontoparietal cognitive control network was to other networks in the brain the less depression symptoms were reported by participants. These results contribute to our understanding of how brain network interactions are related to mental health symptoms, even in individuals who have not been diagnosed with a disorder.

## INTRODUCTION

People vary in their degree of mental health. Indeed, people who do not meet formal criteria for mental illness as defined by our current diagnostic systems (American Psychiatric Association, 2013; World Health Organization, 1992) may still experience a number of symptoms associated with that disorder (World Health Organization, 2017). Here we use this natural variability to better understand neural factors potentially contributing to day-to-day experiences of poor mental health, as well as (prodromal) factors that may elevate the risk for severe mental illness. We hypothesized that the variability observed in mental health symptoms among individuals is related to the function of the frontoparietal cognitive control network (FPN), based on our previously-developed theoretical proposal that the FPN plays a domain-general protective role against mental health symptoms (Cole et al., 2014).

The proposed theoretical framework suggests that alterations in FPN function may play a common role in multiple mental disorders by disrupting a domain-general cognitive control feedback mechanism that can regulate symptoms when they are experienced (Cole et al., 2014). The FPN is a candidate network for this function because it is a flexible hub, meaning it has a high degree of connectivity across the brain (Cole et al., 2010; Power et al., 2011) and can rapidly modify functional connections according to current goals (Cole et al., 2013). There is strong evidence that these FPN functions are domain general (Chein and Schneider, 2005; Cole et al., 2013; Dosenbach et al., 2007), such that individual differences in the general ability to regulate cognition can influence symptoms. Finally, alterations in FPN functional connectivity (FC) have been identified in a number of mental disorders including: depression Kaiser et al., 2015), anxiety (Sylvester et al., 2012), schizophrenia (Baker et al., 2014; Cole et al., 2011; Fornito et al., 2011; Yang et al., 2016), attention deficit hyperactivity disorder (Li et al., 2014; Park et al., 2016), and eating disorders (Boehm et al., 2014; Cowdrey et al., 2014). Consistent with most of these studies, we focus here on FC measured using functional magnetic resonance imaging (fMRI), calculated as the temporal relationship in the blood oxygenation level dependent (BOLD) signal between brain regions (Biswal et al., 1995) while participants rest in the scanner.

Consistent with the flexible hub framework, a number of studies using different measures have provided converging evidence that the FPN is especially well connected to the rest of the brain (Buckner et al., 2009; Cole et al., 2010).

Both of these studies calculated a summary statistic reflecting the degree of connectivity across the whole brain. However, these estimates can be influenced by the relative size of different networks. For example, nodes of a larger network will have a larger overall number of strong connections than nodes of a smaller network simply because, by definition, within-network connections are stronger on average than between-network connections (Power et al., 2013; Wig et al., 2011). Therefore, we estimated how well each region of the brain was connected to the rest of the brain using between-network global connectivity (BGC) (Ito et al., 2017), a measure not influenced by network size.

Particularly important for our specific test of the flexible hub framework here, patients diagnosed with major depression exhibit differences in FC patterns throughout the brain, including FPN functional connections (Brakowski et al., 2017). Specifically, connectivity between regions of the FPN is decreased in depressed individuals (Alexopoulos et al., 2012), as well as in undiagnosed individuals experiencing depression symptoms (Wei et al., 2014). However, Wei and colleagues (2014) looked at FC with specific seed regions, not global connectivity, in their sample. Another study found that global brain connectivity was decreased in the medial prefrontal cortex and the dorsolateral prefrontal cortex (dlPFC) portions of the FPN in depressed patients (Murrough et al., 2016). Depression has been shown to generally disrupt FC in the dlPFC as well as the default mode network (DMN) (for review see: Gong and He, 2015). Decreases in within-network FPN connectivity have also been observed in a group of individuals reporting depression symptoms in the absence of a clinical diagnosis (Hwang et al., 2015). Researchers have also attempted to subdivide depression into various types based on FC patterns. Decreases in FPN connectivity were most pronounced in one particular subtype of depression associated with increased symptoms of fatigue and decreased symptoms of anxiety (Drysdale et al., 2016). Furthermore, activation in the FPN and changes in FPN FC have been observed when participants are actively attempting to regulate their emotions (Banks et al., 2007; Buhle et al., 2014; Kohn et al., 2014) and when participants are in threatening situations (Balderston et al., 2017). Decreased global brain connectivity, the mean FC value for each region’s connections, has been observed in the prefrontal cortex in depression patients (Abdallah et al., 2016). This decrease in global brain connectivity was rescued 24 hours following ketamine treatment. These previous results are broadly consistent with our hypothesis, yet the extension of results to test whether FPN BGC is related to mental health symptoms among healthy individuals would provide important new evidence for the general nature of FPN’s role in regulating mental health.

Consistent with our previously-developed theoretical framework (Cole et al., 2014) along with observed FPN FC alterations in patients with major depression, we hypothesized that individual differences in depression symptoms in undiagnosed individuals would be correlated with BGC in the FPN. Support for our hypothesis would provide important evidence for a potentially general role of global FPN intrinsic FC in facilitating the regulation of mental health symptoms.

## METHODS

### Participants

Data were collected at the Rutgers University Brain Imaging Center (RUBIC). The participants were recruited from the Rutgers University-Newark campus and surrounding community. All participants provided informed consent and all procedures were approved by the Rutgers University-Newark Institutional Review Board. We collected data from 106 participants. Technical error or equipment malfunction during the scanning session resulted in removing six participants from the study. Four participants were removed from the study because they did not complete the Center for Epidemiological Studies Depression Scale (CESD) during a behavior-only session separate from the MRI session. We also collected demographic information and asked participants 11 questions asking what hand they used for various activities including writing, throwing, using a scissors, holding their toothbrush, striking a match, opening a box, kicking, using a knife, using a spoon, which hand was placed on top while using a broom, and which eye they used in situations where they would only being using one eye. They replied to each question by answering always right, usually right, no preference, usually left, or always left. Answers were scored in the following manner: always right (2), usually right (1), no preference (0), usually left (-1), always left (-2). We calculated a laterality quotient (LQ) by summing these scores and dividing by the maximum score of 22. A LQ score of -100 indicates a strong left hand preference, 0 indicates no hand preference, and 100 indicates a strong right hand preference. All participants self reported as right handed. However, one participant demonstrated a LQ indicating they had a left hand preference. The participant with a left hand preference has been included in all analyses.

Studies have proposed that the CESD can be broken down into between one and four factors (Cole et al., 2004; Herrero and Meneses, 2006). In addition to calculating the raw CESD score for each participant we also calculated three factor scores: somatic symptoms, negative affect, and anhedonia, based on a recent study (Carleton et al., 2013). Participants also completed several measures of flexible cognition during the behavior-only session. These measures included Raven’s progressive matrices (Bilker et al., 2012), Cattell’s culture fair test (Cattell and Horn, 1978), and Duncan’s goal neglect task (Dumontheil et al., 2011). The final sample consisted of 96 participants (See Table 1).

**Table 1.** Demographic information

|  |  |
| --- | --- |
| **N** | 96 |
| **Age** | $M = 22.06, SD = 3.84$ |
| **Gender** |  |
| Male | 42 (43.8%) |
| Female | 54 (56.2%) |
| **Handedness (LQ)** | $M = 74.67, SD = 25.2$ |
| **Education (Highest level)** |  |
| High school | 14 (14.6%) |
| Some college | 50 (52.1%) |
| Bachelor's degree | 29 (30.2%) |
| Graduate degree | 3 (3.1%) |
| **Cognitive measures** | (Percent correct) |
| Raven | $M = 52.29, SD = 15.69$ |
| Cattell | $M = 55.20, SD = 12.03$ |
| Duncan | $M = 76.19, SD = 18.34$ |
| **CESD (raw score)** | $M = 17.41, SD = 9.26$ |

### Gender

Multiband whole-brain echo-planar imaging (EPI) acquisition was collected using a 32-channel head coil on a 3T Siemens Trio MRI scanner with the following parameters: TR = 785 ms, TE = 34.8 ms, flip angle = 55°, Bandwidth 1924/Hz/Px, in-plane FoV read = 208 mm, 72 slices, 2.0 mm isotropic voxels, with a multiband acceleration factor of 8. Whole-brain high-resolution T1-weighted and T2-weighted anatomical scans with 0.8 mm isotropic voxels were also collected. Spin echo field maps were collected in both the anterior to posterior direction and the posterior to anterior direction consistent with the Human Connectome Project preprocessing pipelines (Glasser et al., 2013). The resting-state fMRI scan was 14 minutes (1070 TRs) in duration.

### fMRI Preprocessing

Functional imaging data were preprocessed using the Human Connectome Project minimal preprocessing pipeline version 3.5.0. Preprocessing consisted of anatomical restructuring and segmentation, EPI reconstruction, segmentation, and spatial normalization to a standard template, intensity normalization, and motion correction (Glasser et al., 2013). All further processing was conducted in CIFTI 64k greyordinate standard space. The data were parcellated into 360 regions described previously (Glasser et al., 2016), taking the average time series of the vertices within a parcel. At this point all subsequent data analysis was conducted in MATLAB R2014b (The Mathworks). We performed nuisance regression using 12 motion parameters, ventricle and white matter timeseries (as well as their first derivatives), and global signal. We also performed motion scrubbing (Power et al., 2012) based on framewise displacement. Framewise displacement was calculated as the amount of head movement for each frame relative to the previous in terms of Euclidean distance. We next applied a low pass temporal filter (0.3 Hz) to the framewise displacement vector in order to reduce the effect of respiration on our framewise displacement measure (Siegel et al., 2016). The framewise displacement threshold for motion scrubbing was set at 0.3 mm. Motion scrubbing consisted of removing the flagged frame from the timeseries as well as one frame prior and the two frames following. FC was estimated by calculating the Pearson correlation of the BOLD timeseries between each pair of the parcels defined by Glasser and colleagues (Glasser et al., 2016).

### Network Assignment and Analysis

The network assignment of each of the parcels was completed on an independent dataset, the Human Connectome Project (100 unrelated) (Van Essen et al., 2013). Briefly, each parcel was assigned to a network using the Generalized Louvain method for community detection. This process was conducted using resting-state data. We identified 12 functional networks (Spronk et al., 2017) (See Figure 1). The functional network topology findings replicate the major features of several previously published network partitions (Gordon et al., 2016; Power et al., 2011; Yeo et al., 2011).

**Figure 1.**
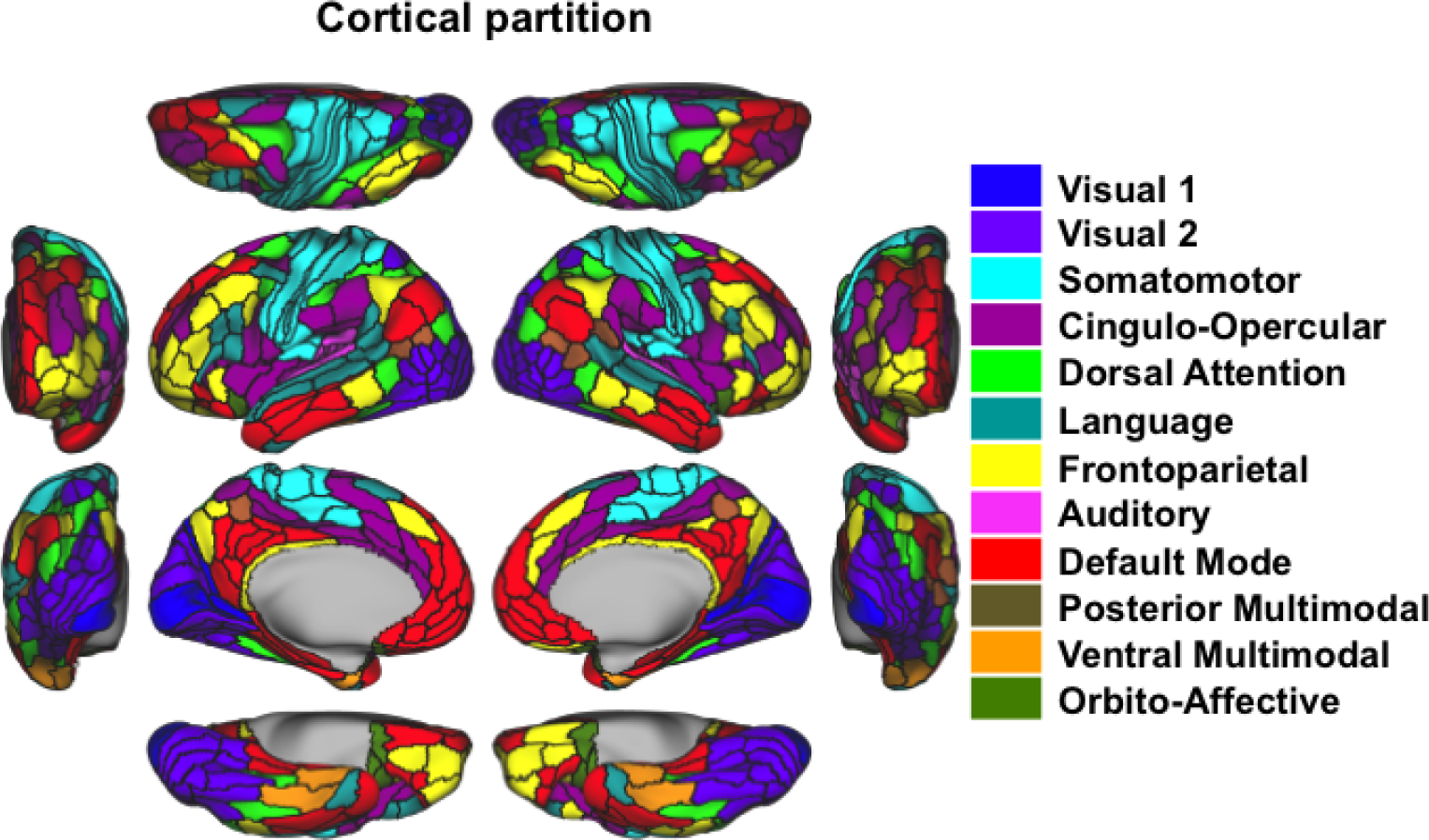
Network assignment. Resting-state fMRI data from an independent dataset (HCP: 100 unrelated) was used to assign each parcel to a network using a community detection algorithm. This resulted in 12 total networks. Color indicates the network assignment for each parcel.

We were interested in the relationship between the frequency at which participants experienced depression symptoms and the degree of between-network global connectivity (BGC). Specifically, we were interested in a measure that would estimate the strength of FC for each region of a network to all of the other regions in different networks. BGC was calculated for each region individually and defined as the mean FC for all out-of-network connections. Out-of-network connections were defined as all connections from a source region to target regions outside the source region’s network. This process was completed for all regions until we had a BGC value for each region in the brain. Then we calculated the mean BGC value for each of the functional networks to summarize effects at the network level.

More formally, BGC was defined for each region as:

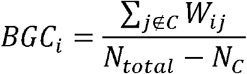

where *BGC*_*i*_ corresponds to the out-of-network weighted degree of region *i* in network *C, j* 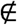 *C* corresponds to all regions not in network *C*, *W*_*ij*_ corresponds to the FC estimate between regions *i* and *j*, *N*_*total*_ corresponds to the total number of regions, and *N*_*C*_ corresponds to the total number of regions in network *C*. We tested the relationship between BGC and depression symptoms in all brain networks. Due to the large number of statistical tests we report the false-discovery rate (FDR)-corrected *p*-values for primary analyses (Benjamini and Hochberg, 1995). Our main hypothesis was that BGC in the FPN was correlated with depression symptoms. Because this was a primary hypothesis based on our previous publication (Cole et al., 2014) we report the uncorrected p-value. Follow-up and control analyses were not independent of the primary tests and are therefore not corrected.

## RESULTS

### Depression Symptom Scores

We hypothesized that the frequency of experiencing depression symptoms is related to individual differences in the between-network global connectivity of the FPN. This would be consistent with the involvement of FPN in domain-general cognitive regulation, broadly construed to include emotion (Cole et al., 2014). We used a standard measure of depression symptoms, the CESD, to measure the frequency of depression symptoms. The CESD consists of 20 questions asking how frequently participants have experienced symptoms related to depression over the last seven days. CESD scores (*M = 17.41, SD = 9.26*) varied from a minimum of 0 to a maximum of 43 (out of a possible 60) (Figure 2). CESD scores from 16 to 24 indicate mild levels of depression symptoms while scores above 24 indicate moderate to severe levels of depression. In our sample 48.9% of participants demonstrated at least mild levels of depression, while 25% of participants exhibited at least moderate levels of depression. The characteristics of our sample were consistent with previous studies using the CESD with undiagnosed young adults (Gress-Smith et al., 2015; Van Dam and Earleywine, 2011). However, the scores were not normally distributed based on a Kolomogorov–Smirnov test (*p < 0.023*). We therefore used the Box-Cox power transformation (Box and Cox, 1964), a common approach to correct for non-normality. After the transformation the data no longer deviated from a normal distribution (*p = 0.17*). The transformed data were used for all subsequent analyses. CESD scores were not correlated with any of the flexible cognition measures (*smallest p = 0.28*).

**Figure 2.**
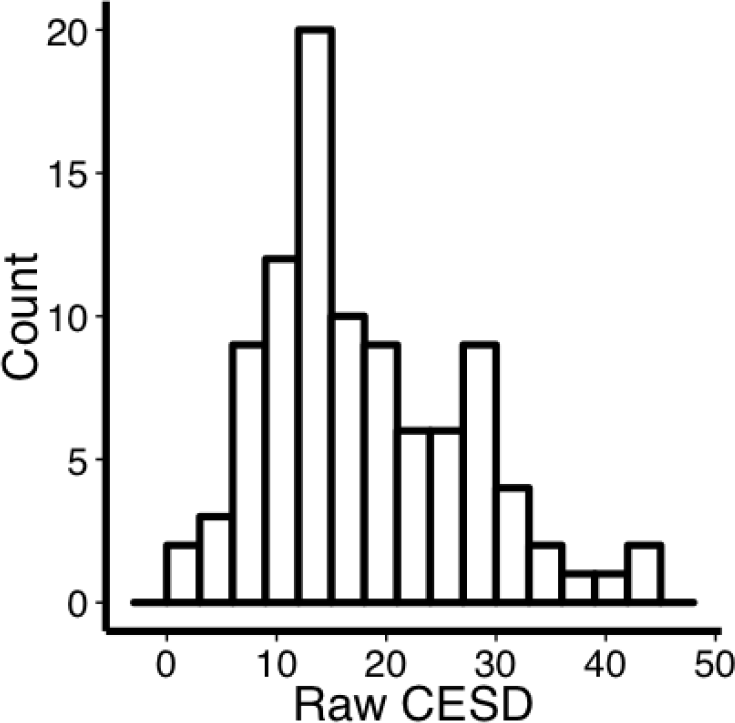
Histogram of raw CESD scores.

We also calculated CESD sub-scores for somatic, negative affect, and anhedonia factors. After Box-Cox transformation none of the factor score distributions significantly deviated from a normal distribution (*somatic p = 0.4, negative affect p = 0.19, anhedonia p = 0.056*). All three of the factors were significantly correlated with the overall CESD scores (*somatic r = 0.77, uncorrected p < 0.001; negative affect r = 0.80, uncorrected p< 0.001; anhedonia r = 0.52, uncorrected p < 0.001*, See Table 2). The three factors were also correlated with each other (*correlation between somatic and negative affect r = 0.55, correlation between somatic and anhedonia r = 0.18, correlation between negative affect and anhedonia r = 0.26*, See Table 1). Although all of the factors were highly related to the overall CESD score, they appear to have contributed unique variance as the highest correlation between any two factors was only accounting for 30% of the variance (based on R^2^ values).

**Table 2.** Correlation between the CESD and CESD factors

|  | Overall CESD | Somatic | Negative<br>Affect | Anhedonia |
| --- | --- | --- | --- | --- |
| Overall CESD | ----- | ----- | ----- | ----- |
| Somatic | 0.77 | ----- | ----- | ----- |
| Neg. Affect | 0.80 | 0.55 | ----- | ----- |
| Anhedonia | 0.52 | 0.18 | 0.26 | ----- |

### Between-Network Global Connectivity

We calculated BGC values using resting-state fMRI data for each functionally defined cortical parcel (Figure 3). DMN regions tended to have lower BGC values, consistent with the conceptually-similar participation coefficient results reported by Power et al. (2011). In contrast, the FPN had high levels of BGC in lateral prefrontal portions of the network (*M = 0.002, SD = 0.02*) relative to the mean of regions outside of the FPN (*M = -0.0081, SD = 0.013; t(95) = 5.08, p < 0.0001*). However, FPN had lower BGC values in parietal (*M = -0.0258, SD = 0.0190*), inferior frontal (*M = -0.0177, SD = 0.0209*) and temporal lobe regions of the network (*M = -0.0420, SD = 0.0221*) relative to the mean of regions outside of the FPN (*largest p < 0.0001*). Sensory networks exhibited two patterns. The visual network had some of the lowest BGC values in the brain with the primary visual cortex region showing the least BGC in the network (*M = - 0.0742, SD = 0.0348*). In contrast, the auditory network (*M = 0.0212, SD = 0.0171*) and portions of the sensorimotor network surrounding the central sulcus (*M = 0.0117, SD = 0.0249*) both showed a high degree of BGC. The sensorimotor regions closest to the central sulcus showed more moderate BGC scores (*M = -0.0139, SD = 0.0289*).

**Figure 3.**
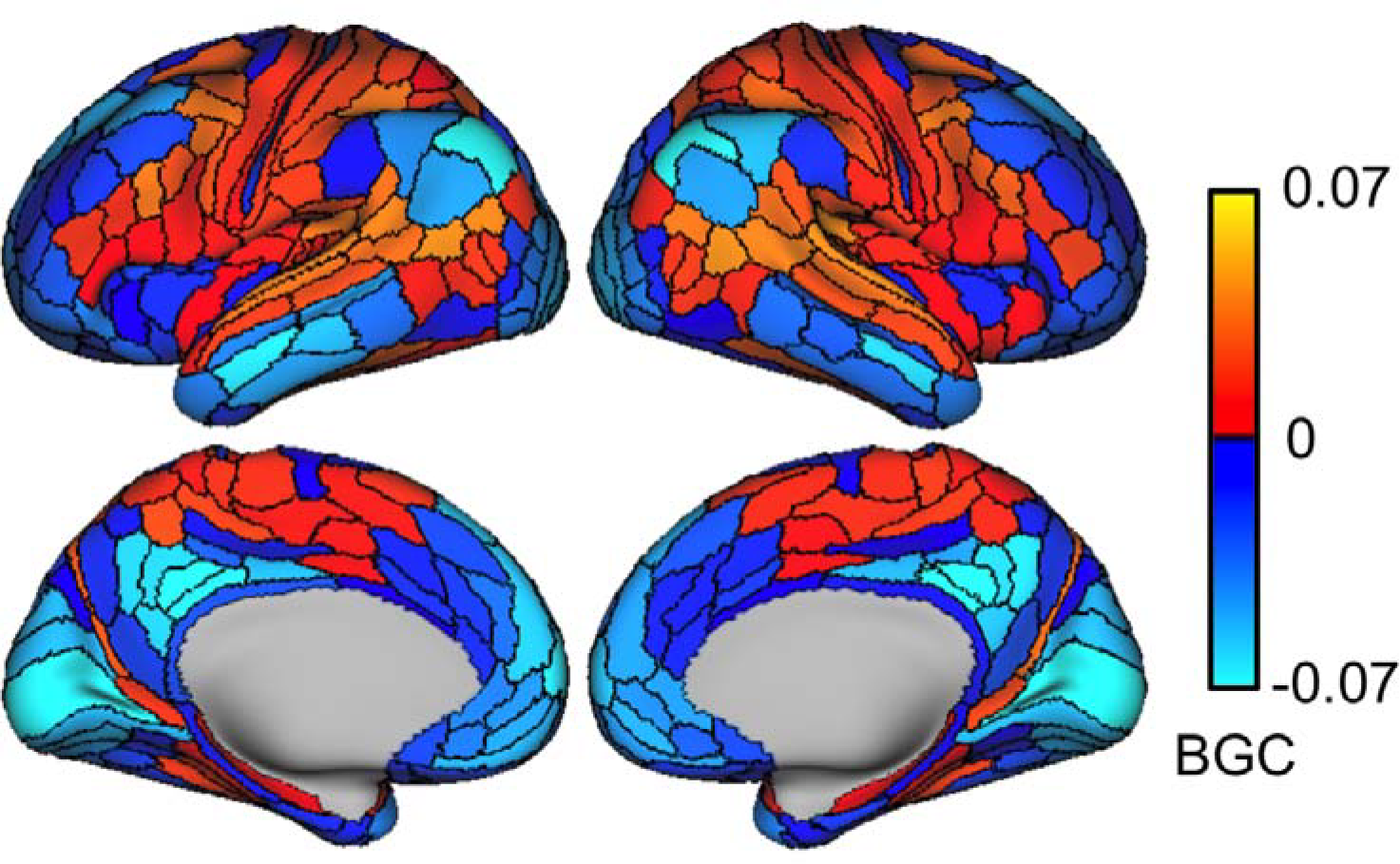
Between network global connectivity (BGC) across all cortical regions. Between network global connectivity for region A is defined as the mean FC (Pearson correlation) between that region A and all other regions outside of region A’s network. Warm values indicate positive values and cool values indicate negative values.

### Correlation Between BGC and Depression Symptoms

We next tested for a relationship between BGC and depression symptoms. This involved creating a mean BGC score for each functional network and testing for correlation between those values and the depression symptom measure (See Table 3). Consistent with our a priori hypothesis, we found that FPN BGC was significantly correlated with depression symptoms (*r* = *-0.247, p = 0.015*) (Figure 4A). Unexpectedly, we found that DMN and language network BGC showed a similar magnitude effect (DMN *r = -0.241, uncorrected p = 0.018;* language *r = -0.268, uncorrected p = 0.008*) (Figure 4B & 4C). These findings suggest that greater FC between the FPN and the rest of the brain (outside of the FPN) is related to less frequent depression symptoms. Less depression symptoms are also related to greater BGC in the DMN and language networks.

**Figure 4.**
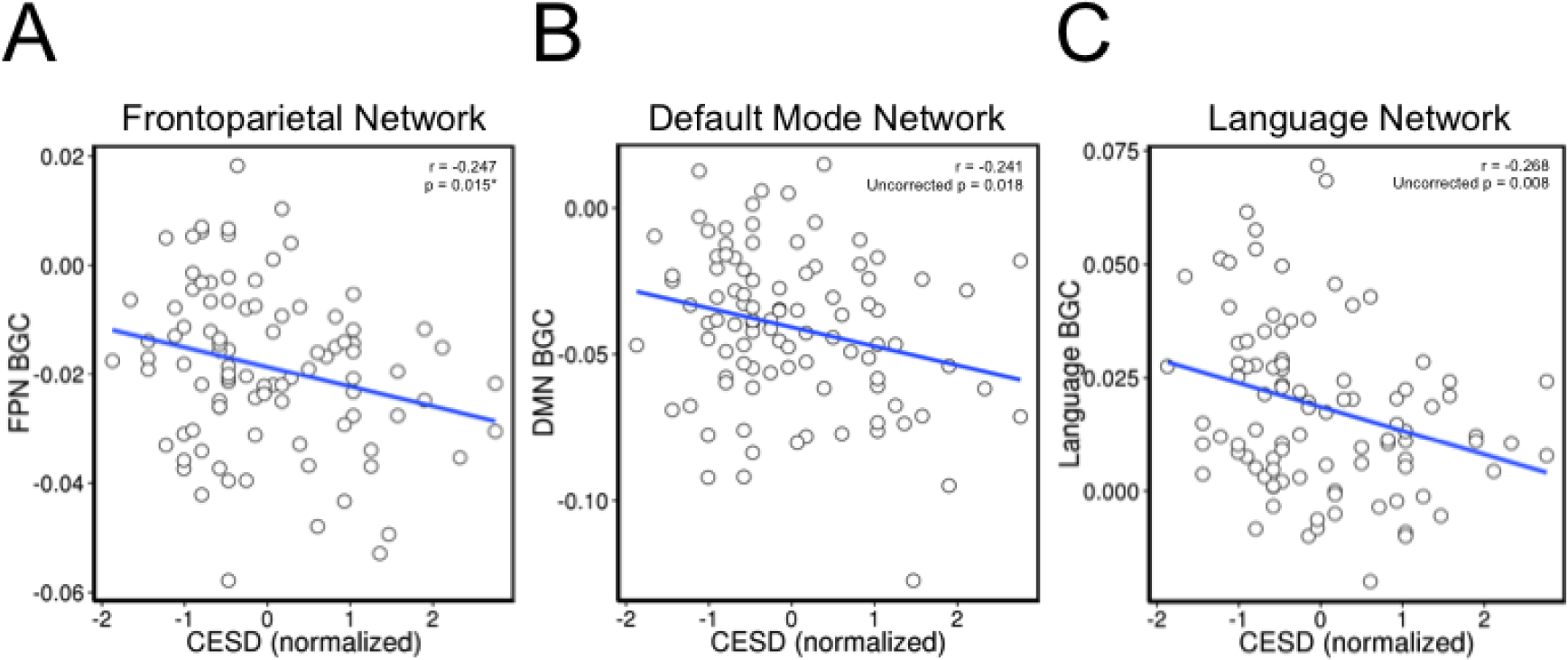
CESD scores and BGC in the FPN, DMN, and language networks are negatively correlated. A) BGC in the FPN is plotted on the y-axis and CESD scores are plotted on the x-axis. B) BGC in the DMN is plotted on the y-axis and CESD scores are plotted on the x-axis. C) BGC in the language network is plotted on the y-axis and CESD scores are plotted on the x-axis.

**Table 3.** Correlation between BGC and CESD scores

| Network | Pearson's r | uncorrected p-value |
| --- | --- | --- |
| Visual 1 | 0.015 | 0.884 |
| Visual 2 | 0.020 | 0.850 |
| Somatomotor | -0.177 | 0.085 |
| Cingulo-Opercular | -0.143 | 0.164 |
| Language | -0.268 | 0.008 |
| Default Mode | -0.241 | 0.018 |
| **Frontoparietal** | **-0.247** | **0.015\*** |
| Auditory | -0.099 | 0.340 |
| Post. Multimodal | -0.110 | 0.284 |
| Dorsal Attention | -0.113 | 0.275 |
| Ventral Multimodal | 0.116 | 0.261 |
| Orbito-Affective | -0.078 | 0.448 |

We report an uncorrected p-value for the test of our a priori hypothesis that FPN BGC would be related to depression symptoms. We also examined the relationship between depression symptoms and BGC in the other 11 brain networks. These p-values do not survive FDR correction for multiple comparisons, but because the magnitude of the effect observed in the DMN and language network was similar to that in the FPN we continue to report subsequent uncorrected statistics for these networks. Although it should be noted that the DMN and language network results should be interpreted with caution as they were not a priori hypotheses and do not survive multiple comparison correction.

To address whether these findings were specific to this particular network partition we conducted a similar analysis using a different network partition scheme (Ito et al., 2017). This network partition includes a FPN and DMN, but lacks a network analogous to the language network. Using this network partition we found a similar negative correlation between BGC and depression symptoms in the FPN (*r* = *-0.262, p = 0.009, FDR adjusted for multiple comparisons p = 0.044*) and DMN (*r = -0.250, p = 0.014, FDR adjusted p = 0.049*). The results for the FPN and DMN both survive correction for multiple comparisons with this network partition.

We conducted a similar analysis, correlating BGC and depression symptoms on the region level, to examine if there were particular brain regions within these networks driving the network level results. None of the individual brain regions showed a significant correlation after correcting for multiple comparisons using FDR. This suggests that the network level results are not being driven by particularly strong relationships in smaller number of brain regions within these networks.

In order to verify that these correlations were not dependent on the Box-Cox transformation we used to normalize the depression symptom data we also ran non-parametric Spearman’s rank correlations, which do not require normally distributed data. The results of testing our a priori hypothesis that BGC in the FPN would be related to depression symptoms were unchanged. Specifically, the BGC-depression rank correlation was consistent with significant Pearson correlations (FPN *rho = -0.227, p = 0.026*). The Spearman rank correlations also reflected similar results to the Pearson correlation in the DMN (*rho = -0.220, uncorrected p = 0.031*) and the language network (*rho = -0.285, uncorrected p = 0.005*).

To ensure that our observed correlation between depression symptoms and BGC in the FPN was not being driven by other factors we removed the variance in the CESD scores that could be accounted for by age and gender factors with a regression model and used the residuals to rerun the correlation. We still found a significant relationship between BGC in the FPN and depression symptoms (*r = -0.220, p = 0.031*). This result suggests that the relationship between BGC in the FPN and depression scores is not being influenced by age or gender differences in our sample.

Next, we tested if the relationships between BGC and overall depression scores were driven by specific depression factors. We tested for correlation between the BGC values and three factors derived from the CESD questionnaire: somatic symptoms, negative affect, and anhedonia symptoms (Carleton et al., 2013). The three factors were correlated with one another so we also used a partial correlation approach to explore the relationship between the unique variance contributed by each factor and BGC in the FPN, DMN and language networks. The partial correlation approach allows us to examine the relationship between BGC and each of the factors while accounting for the variance contributed by the other factors. After accounting for the variance contributed by other factors we found that none of the factors were significantly correlated with BGC in the FPN (*largest rho = -0.110, p = 0.292*). We also found that the unique variance from each factor was not correlated with BGC in the DMN (*largest rho = -0.137, p = 0.188*). BGC in the language network was not significantly correlated with the unique variance contributed by any of the factors (*largest rho = -0.185, p = 0.075*). These results suggest that the relationships we observed between the overall CESD scores and BGC were driven by a combination of all factors, and not specific features of depression.

### Observed BGC-depression Correlations were not Dependent on Connections Between FPN, DMN, and language network

The magnitude and direction of the BGC-depression correlations were similar for the FPN, DMN, and language network. One possible explanation for these results is that the observed BGC-depression effects for these networks were driven primarily by the FC between them. We tested this possibility by recalculating BGC for each network by removing connections between the FPN, DMN, and language network from the calculation. This approach minimizes the possibility that connections between these networks drove the original correlations between depression symptoms and BGC The magnitude and direction of the correlations between depression symptoms and BGC in the FPN (*r = -0.226, p = 0.027*), the DMN (*r = -0.236, uncorrected p = 0.021*), and the language network (*r = -0.201, uncorrected p = 0.049*) were similar when we excluded the connections between these networks from the BGC calculation.

This suggests the original BGC-depression correlations were not dependent on connections between the FPN, DMN, and language network.

In order to further explore the possibility that the observed relationship between depression symptoms and BGC in the FPN might be primarily driven by FPN connections to the DMN and language network we calculated the correlation between the mean FPN to DMN, and FPN to language network FC with depression symptoms. There was not a significant correlation between depression symptoms and the mean connectivity between the FPN and DMN (*r = -0.016, uncorrected p = 0.881*) or the mean connectivity between the FPN and language network (*r = -0.146, uncorrected p = 0.156*). This suggests that the relationship between FPN BGC and depression symptoms is not driven by connections to the DMN and language networks, which both showed a negative correlation between BGC and depression symptoms as well.

### Within-Network Connectivity does not Correlate with Depression Symptoms

We focused on between-network connectivity, yet the degree of connectivity within a functional network might also be related to depression symptoms. For example, a high degree of FC within a network may reflect more homogeneous network activity. In this case the network would be tightly coupled and all of the component regions would likely be serving a very similar function. However, a lower degree of FC within a network may reflect more heterogeneous processing. In this case the different regions of a functional network may all be involved in the same general process, but they may be contributing in different ways. These differences in within-network functionality may, in turn, relate to depression symptoms.

Consistent with our choice to focus primarily on between-network effects, within-network FC in the FPN was not significantly correlated with depression symptoms (*r = 0.066, p = 0.526*). Within-network FC in the DMN was also not significantly correlated with depression symptoms (*r = 0.143, uncorrected p = 0.164*). The language network showed a negative correlation between depression symptoms and within-network FC (*r = -0.219, uncorrected p = 0.032*). We observed a significant difference in the DMN correlation coefficients comparing BGC (*r = -0.241*) and within-network connectivity (*r = 0.143*) with depression symptoms (*z = 2.66, p = 0.008*). It will be important for future studies to test this effect for replication, given the non-significance of the DMN within-network FC-depression effect relative to a correlation of 0.

## DISCUSSION

Based on convergent evidence across a variety of mental health conditions we predicted that individual differences in FPN BGC would be correlated with symptoms associated with depression (Cole et al., 2014). This is consistent with extensive evidence that the FPN is a domain-general cognitive control system regulating general goal pursuit processes (Cole and Schneider, 2007; Duncan, 2010; Schneider, 2003), including regulation of mental illness symptoms (Cole et al., 2014). We sought to expand the general relevance of this framework by including symptoms experienced in everyday life, which express dimensionally across health and disease. This involved investigating the relationship between the mood symptoms, one of the most commonly experienced set of symptoms in the general population (Centers for Disease Control and Prevention (CDC), 2010), and FPN global connectivity properties. We further ensured the broad relevance of the findings by focusing on how a general population who had not been diagnosed with a mental disorder varied in their depression symptoms and primarily experienced them in the normal (i.e., sub-clinical) range.

As predicted, we found a significant relationship between how well-connected the FPN was to the rest of the brain and the frequency of depression symptoms in adults who had not been diagnosed with depression. We used BGC to estimate the degree of brain-wide connectivity for each brain region. The results suggest that individuals who report fewer depression symptoms are characterized by a FPN that is more connected to the rest of the brain. These results support our previously developed theoretical framework, suggesting natural variance in FPN global connectivity influences each individual’s ability to regulate mood symptoms in everyday life. Extending this framework to test if FPN function relates to natural variation in other symptom domains will be critical as well as examining if such effects persist when an individual crosses a threshold necessary for a formal diagnosis.

### Between-network global connectivity identifies how well each brain region is connected to other brain networks

We hypothesized that individuals exhibiting greater FC between the FPN and the rest of the brain would report fewer depression symptoms. We used BGC, a measure that calculates the mean FC between each brain region and all other out-of-network brain regions (Ito et al., 2017), to evaluate this hypothesis. BGC reduces the potential bias of other graph centrality measures, which can be inflated in regions assigned to large networks (Power et al., 2013). Previous methods using global brain connectivity and degree, which both include within network connections, result in DMN regions showing high connectivity with the rest of the brain (Buckner et al., 2009; Cole et al., 2010; Liang et al., 2013).

However, we found that BGC was relatively low in the DMN. This is consistent with DMN results from studies using participation coefficient (Power et al., 2013, 2011). While our measure of BGC showed similarity to results obtained using participation coefficient (Power et al., 2011), there were some differences. Some of these differences may be due to the differences between BGC and participation coefficient. Participation coefficient is generally calculated by first thresholding a graph and then examining the distribution of edges between networks. A high value indicates that the edges are relatively evenly distributed to other networks and lower values indicate that the edges tend to preferentially connect nodes in a smaller number of other networks. BGC does not require thresholding a graph prior to calculation and simply measures the mean FC weight for each region to all other out of network regions. These results suggest that previous findings identifying the DMN as highly connected to the rest of the brain are largely driven by high FC within the DMN, and do not necessarily reflect greater connectivity between the DMN and nodes in other functional networks.

The results we observed with BGC were similar to previous attempts to identify hubs and the most well connected regions in the brain. BGC was high in the lateral prefrontal cortex, the motor and tactile cortex, the auditory cortex, and higher order visual regions. Lateral prefrontal regions and higher order visual areas show a greater degree of FC (Buckner et al., 2009) and a higher global brain connectivity (Cole et al., 2010). Higher global brain connectivity has also been reported in the lateral prefrontal cortex, higher order visual regions, auditory cortex, and somatosensory cortex (Liang et al., 2013). We observed that BGC was consistent with previous attempts to classify the degree of connectivity in many brain regions.

There were some discrepancies between previous methods and BGC. We found that BGC was quite low in lower visual regions in contrast to higher connectivity estimates calculated by others (Cole et al., 2010; Liang et al., 2013). The differences observed between BGC and other measures of connectivity strength may be driven by BGC not considering the relatively strong local connections within the visual network. In fact, primary and secondary visual cortex show high local connectivity relative to distant connectivity strength (Sepulcre et al., 2010). It will be important for future studies to identify whether differences observed in primary sensory cortices are driven by an imbalance between within and between-network connectivity.

### BGC in the FPN is negatively correlated with depression symptoms

We found that BGC in the FPN showed a significant negative correlation with depression symptoms. This suggests that individuals exhibiting greater connectivity of the FPN with the rest of the brain experience fewer symptoms of depression. The correlation between BGC in the FPN and depression symptoms supports our hypothesis that a well-connected FPN may serve a protective role against depression symptoms and possibly mental health symptoms in general (Cole et al., 2014).

Decreases in FPN FC have been reported in individuals diagnosed with major depression (Alexopoulos et al., 2012; Murrough et al., 2016; Veer, 2010). Differences in FC patterns can also be used to divide depression into distinct subtypes (Drysdale et al., 2016). We build on these findings by observing similar results in a variable sample of undiagnosed individuals that likely includes primarily mentally healthy, but also some mentally unhealthy, participants.

Another study examining FC in a group of undiagnosed individuals with higher levels of depression symptoms found that connectivity between the superior parietal lobule and the dlPFC portion of the FPN was decreased (Wei et al., 2014). Undiagnosed participants experiencing more depression symptoms have also been reported to have reduced FC between dlPFC and the supramarginal gyrus, insula, operculum, precuneus and parahippocampal gyrus (Hwang et al., 2015). Previous studies have also found that anti-depressant and electroconvulsive therapy treatments can modulate prefrontal cortex FC in depression patients (Perrin et al., 2012; Wang et al., 2015). It is possible that future therapeutic interventions could target and attempt to alter aspects of network communication. Our findings suggest that greater global connectivity of the FPN is related to reduced depression symptoms. Our data regarding global connectivity differences is consistent with a recent study that found reduced global brain connectivity in the lateral PFC in depression patients (Abdallah et al., 2016). Here we extend the relationship between global connectivity in the FPN to include samples that have not been diagnosed with a clinical disorder and further refine the global connectivity findings by suggesting that the between network connections are particularly important. Another recent study has suggested that the FPN can be split into two subsystems, one more involved in introceptive processes, and another more involved in the regulation of perceptual attention (Dixon et al., 2018). Future studies should examine if FC relationships with depression are stronger in the introceptive FPN system that may be more involved in rumination symptoms.

Unexpectedly, we also found a negative correlation between BGC in the DMN and depression symptoms. This result did not survive multiple comparison correction, but because it was of a similar magnitude as our predicted relationship in the FPN we have explored it. It is important to note that the results in the DMN should be interpreted with caution and future studies should attempt to replicate these results. DMN activity has been linked to rumination symptoms in depression (Hamilton et al., 2011). This finding may reflect an inability for individuals experiencing more depression symptoms to disengage the DMN in situations when attentional or cognitive resources need to be allocated (Sheline et al., 2009). Depression patients also exhibit decreased FC between the DMN and executive networks (Abbott et al., 2013; Manoliu et al., 2014). These findings provide further support for the interpretation that depressed individuals may have a difficult time disengaging the DMN because the connections from cognitive control networks to the DMN are decreased.

A number of studies have suggested that FC within portions of the DMN is increased in depression patients (Greicius et al., 2007; Li et al., 2013). Although we did not find a significant correlation between within-network FC in the DMN and depression symptoms, we did see a trend toward a positive correlation. Our lack of a correlation between within-network FC in the DMN and depression symptoms could be due to methodological differences in using a population of individuals who have not been diagnosed with a disorder versus comparing a control group to a group diagnosed with depression. Additionally, we considered the entire DMN rather than using a seed correlation approach. It will be important for future studies to investigate the relationship between within-DMN FC and depression symptoms, verifying this effect in a larger sample and tying it to specific neural mechanisms and specific depression symptoms.

We also found a negative correlation between BGC in the language network and depression symptoms. Similar to the DMN, this result did not survive multiple comparison correction, but because the magnitude of the effect was similar to that of our predicted relationship between BGC in the FPN and depression symptoms we have explored it. Because we did not have an a priori prediction about this network and the result did not survive multiple comparison correction, the interpretation of this result should be tempered. Decreases in FC in the language network have been reported in depression patients (Buchanan et al., 2014) and language performance deficits have also been reported in depression patients (Baune et al., 2010). The current results might be consistent with these observed deficits in language in depressed individuals. However, future studies should attempt to replicate these results in an independent large sample.

## Conclusion

The current study sought to test our previously-developed framework that suggests the FPN (along with other cognitive control networks) acts as a protective factor against mental disease via its widespread FC with other networks (Cole et al., 2014). We identified a negative correlation between depression symptoms and a measure of between-network, global FC in the FPN in a sample of individuals from the general population who had not been diagnosed with depression. These results suggest the human brain’s global network architecture is critical for maintaining mental health even in undiagnosed individuals, supporting the possibility that FPN maintains a goal-directed feedback loop to regulate symptoms as they arise. It will be important for future research to characterize the exact mechanisms by which FPN influences symptoms, and to assess the possibility of enhancing FPN FC in the interest of reducing symptoms and potentially preventing the onset of mental illness.

## Acknowledgements

This work was supported by the National Institute of Health (MH096801, MH109520, and AG055556). The content is solely the responsibility of the authors and does not necessarily represent the official views of any of the funding agencies.

